# Social attachment shapes interbrain synchrony

**DOI:** 10.64898/2026.04.15.718291

**Authors:** Kathleen Murphy, Liza E. Brusman, Yevgenia Kozorovitskiy, Zoe R. Donaldson

## Abstract

Neural synchrony, or correlated neural activity across interacting individuals, scales with relationship quality in humans, yet how it evolves during social bond formation remains unknown. Using fiber photometry in monogamous prairie voles, we track prefrontal cortex synchrony across pair bond formation. Bonded voles show stronger synchrony with partners than strangers, mirroring human findings. A linear mixed model reveals that synchrony is jointly shaped by bond strength, inter-animal distance, and time since interaction onset, with relationship type modulating how each factor contributes. Using a machine-learning behavioral classification pipeline we developed for freely interacting voles, we demonstrate that the coupling between specific behaviors and synchrony depends on the nature of the dyadic relationship. These findings establish that neural synchrony is not a simple function of proximity or interaction time but is fundamentally shaped by relationship history—a conclusion with direct implications for understanding the synchrony in human social attachment.

---

While neuroscientists have historically examined single brains in isolation, social behavior emerges not in isolation, but from dynamic interactions between individuals. One measurable signature of these dynamic interactions is interbrain synchrony, the correlated neural activity that emerges between individuals engaged in shared social experience^1–3^. Hyperscanning approaches—using fMRI, EEG, MEG, or fNIRS to record from multiple subjects simultaneously—have revealed that this interindividual neural synchrony reflects more than shared sensory input^4–13^. For example, it scales with relationship quality and social dynamics^14–21^. Romantic partners show greater synchrony than strangers during identical tasks^18^, synchrony magnitude correlates with empathy and cooperative behavior^18,22,23^, and synchronization patterns shift dynamically during interactions^24,25^. Thus, some facets of interbrain synchrony may index the nature of a given relationship.

This coupling between brains is conserved across several mammalian species^26–29^. Recordings of electrical activity and calcium signals in the prefrontal cortex of bats and mice, respectively, demonstrate that correlated neural oscillations between individuals emerge during social interactions and can be detected across recording modalities that vary in spatiotemporal resolution^28,29^. Further, bats show stronger synchrony during vocal communication with members of the same social ‘clique’, while synchrony in mice scales with the disparity in social hierarchy standing^28,29^. Together, these data suggest that prior social experience with a particular individual affects interbrain synchrony.

The dynamics of neural synchrony during relationship formation remain poorly understood, in part because it is rarely possible to study human pairs before and after social bond formation, necessitating the use of another species that displays human-like abilities to form lasting social bonds. Here, we use monogamous prairie voles, a species that readily forms selective, enduring pair bonds, to examine interbrain synchrony before and after bond formation^30–34^. Using photometry-based measurements of prefrontal cortical Ca^2+^ activity, we find that prairie voles, like mice and bats, exhibit elevated interbrain synchrony during social interaction. Bonding leads to stronger interbrain synchrony with a partner compared to a stranger, mirroring human findings. Synchrony is best predicted by a model incorporating the type of relationship between animals, bond strength, distance between animals, and time since interaction initiation. Using a machine-learning-based behavioral classification pipeline we have developed for freely interacting prairie voles, we further find that the relationship between social behavior and interbrain synchrony is itself modulated by the nature of the dyadic relationship. Together, these findings elucidate the sensory, cognitive, and behavioral factors that contribute to interbrain correlation (IBC)—all dynamically shaped by the history, quality, and behavioral context of a relationship.

## Results

### Prairie voles have increased IBC in the mPFC during social interaction

To examine the relationship between IBC and social interaction in monogamous prairie voles, we used fiber photometry recordings of GCaMP-mediated Ca^2+^ activity in the medial prefrontal cortex (mPFC) of opposite-sex pairs before and after pair bond formation (Fig. 1A, B). Sexually naive voles were recorded while interacting sequentially with two different opposite-sex voles, one of which was randomly assigned to become their future partner (Fig. 1A, C). Following initial recordings, voles cohabitated with their assigned partner, and we used a partner preference test to quantify bond strength (Fig. 1D). After two weeks of cohabitation, we repeated sequential IBC recording sessions with their partner and an opposite-sex stranger, with the order randomized.

**Figure 1.**
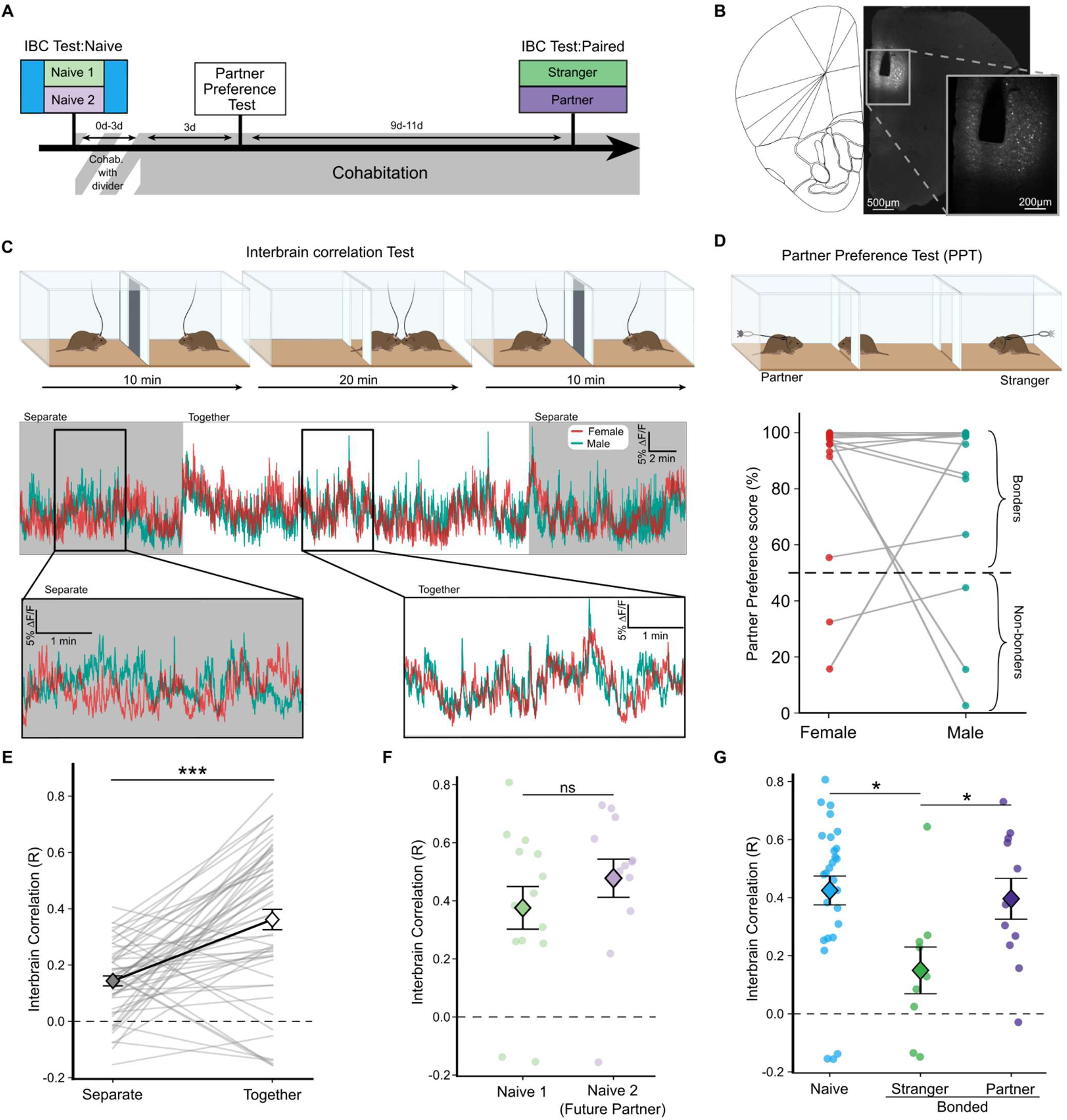
Prairie voles show relationship-dependent increases in interbrain correlation (IBC) during dyadic interaction. **A.** Dyadic recordings of GCaMP fluorescence were carried out with two opposite-sex individuals prior to and following cohabitation. Bond formation was tested via a partner preference test (PPT). **B.** Prairie voles expressing GCaMP6f were implanted with an optic ferrule in the mPFC. **C.** (Top) Interbrain recording sessions consisted of 40-minute simultaneous fiber photometry recordings, which included a 20-minute free interaction portion (together), bookended by 10-minute portions where voles were separated by an opaque divider. (Bottom) Example trace showing normalized fluorescence for male and female voles across the entire IBC test. Example 5-min periods of the above sample trace from the separated (left) and together (right) periods. **D.** (Top) Bond strength was measured using a 3-hr PPT. Most voles formed bonds, evidenced by more time spent huddling with their partner than with a stranger (% partner preference). (Bottom) Voles with less than a 50% partner preference score (n = 5 animals) were considered non-bonders. Lines connect partners. **E.** IBC was stronger during periods of social interaction compared to periods of separation; data from all recordings, paired t-test: t_52_ = −5.90, p = 2.8e^−7^. **F.** Before pairing, there was no significant difference in IBC between two potential future mating partners (Naïve 1 and Naïve 2); independent t-test: t_25_ = −0.98, p=0.336. **G.** After bonding, IBC is stronger between partners than strangers; two-tailed independent t-test, t_16_ = −2.24, p = 0.039. IBC depends on the relationship between voles; ANOVA, F_2,43_ = 4.84, p=0.013. Bonding leads to decreased IBC with strangers compared to naïve interactions (naïve 1 and 2 aggregated, blue); Tukey HSD: Naive and Partners p-adj = 0.983, Naive and Strangers: p-adj = 0.028. All error bars show mean and standard error. *p<0.05, **p<0.01, ***p<0.001

To measure interbrain correlation between voles, we compared IBC calculated using Pearson’s correlation coefficient during session periods when the voles were separated by an opaque divider (separated) and those without a divider, when unconstrained interactions occurred (together). Voles exhibited more correlated mPFC activity during the interaction sessions than when they were separated (Fig 1E; paired t-test t_52_ = −5.90, p = 2.76e^−07^). As both voles were exposed to shared external sensory stimuli (lights, sounds, etc.) but were unable to interact or see each other when separated by a divider, this observation supports at least a partial dissociation of socially-mediated IBC from that attributed to similar responses to the shared environment.

### IBC during free interaction decreases with strangers after bond formation

Next, we investigated how IBC during social interaction differs as a function of relationship type, that is, with another sexually naive opposite sex vole (naïve), a bonded partner (partner), or a stranger encountered after bonding (stranger). We first tested the hypothesis that pair-bonding would lead to stronger IBC with partners than with strangers. Comparing IBC during partner and stranger interaction for pairs where both voles reached a threshold of partner preference >50%, we found that bonded voles have stronger IBC with their partners than with strangers (Fig. 1G; t_16_ = - 2.24, p = 0.039).

Next, we asked whether this difference in partner and stranger-associated IBC reflects relative strengthening or weakening of synchrony, compared to that exhibited towards opposite-sex voles before bonding (*i.e*., with potential mates). Here, we pooled all interactions with potential mates into a single group (naïve) after confirming that there was no significant group-level difference in IBC between the naïve 1 and naïve 2 subgroups (Fig. 1F, ind. t-test, t_25_ = −1.03, p = 0.313). Surprisingly, bond formation did not drive stronger IBC with partners compared to the naive group, but rather decreased IBC with subsequently encountered strangers (Fig. 1G, Supp. Fig. 1A; ANOVA, (F_2, 42_ = 3.72, p=0.033; Tukey HSD: naïve vs partner: p = 0.983; naïve vs stranger: p = 0.028).

### Variation in IBC is not fully explained by bond strength or interaction time alone

To identify factors that contribute to variation in IBC strength, we examined several potentially simple explanations. First, we asked whether IBC during naive interactions predicted future bond strength (average pair partner preference), and whether bond strength predicted IBC between bonded voles at later timepoints (Fig. 2A). We did not observe a significant correlation in either direction (Fig. 2B: naïve IBC → future bond strength: r = 0.32, p = 0.279; Fig. 2C; bond strength → IBC between partners: r = 0.34, p = 0.235). Bond strength also did not predict IBC when interacting with a stranger after bonding (Fig. 2C; r = −0.28, p = 0.381). While these correlations were not significant, they all trended in the expected direction (e.g. a positive slope for bond strength:IBC for partners, negative for strangers), potentially suggesting that the dataset may be underpowered to detect these correlations within segmented subgroups. Alternatively, and consistent with our modeling data described below, detecting the effect of bond strength on IBC may require a multivariate approach to account for the interacting effects of time and behavior on IBC.

**Figure 2.**
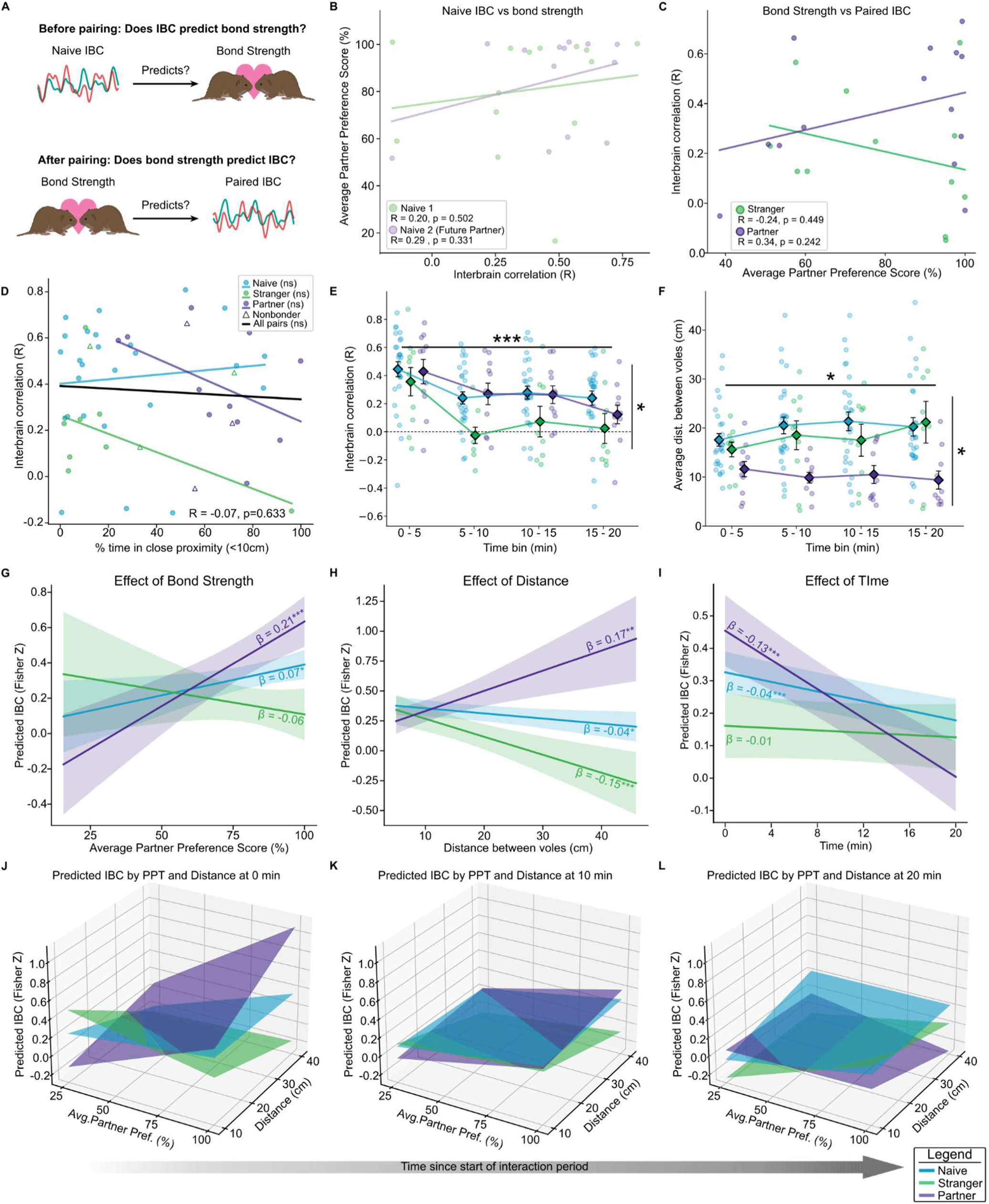
The effect of bond strength and inter-animal distance on IBC depends on relationship type. **A.** We asked whether pre-pairing (naïve) IBC predicts future bond strength and whether bond strength predicts post-pairing IBC. **B-D.** Pearson correlations between IBC and bond strength or IBC and percent time in proximity for any relationship type were not significant (all P > 0.05). **E.** IBC varies across the recording session and by relationship type (mixed ANOVA: time, F_3,123_ = 18.98, p_GG-corr_ = 6.2e-9; relationship, F_2,41_ = 3.54, p = 0.038). IBC is highest during the first 5 min and is greater in naïve and partner dyads than strangers (Holm-corrected and Tukey HSD post hoc tests). **F.** Mean inter-animal distance also varies across time and relationship type (mixed ANOVA: time, F_3, 117_ = 3.52, p = 0.012; relationship, F_2,39_ = 5.00, p = 0.017), with partners remaining closer than naïve or stranger dyads. **G-I.** After bond formation, the effects of bond strength, inter-animal distance, and time in session on IBC diverge by relationship type. Shaded bands, 95% CIs. **G.** Positive association between IBC and bond strength in naïve and partner interactions. **H.** Positive association between IBC and distance in partners but negative associations in naïve and stranger interactions. **I.** IBC declines over the session for all groups, except strangers. **J-L.** The effects of bond strength and inter-animal distance attenuate over time, shown at session start (J), midpoint (K), and end (L). Error bars, mean ± sem. *p < 0.05, **p < 0.01, ***p < 0.001. Shared legend applies to E–L.

We next asked whether pairwise differences in IBC reflect differences in how much a pair interacted during a recording session. We used the proportion of time animals spent within 10 cm of each other (∼1 vole length) as a proxy for social interaction^35,36^. There was no significant correlation between social interaction time and IBC across all recordings (Fig. 2D; R = −0.07, p = 0.633), or within each of our three groups (Fig. 2D; naive: R = 0.10, p = −0.637, partners: R = −0.46, p = 0.154, strangers: R = −0.54, p = 0.171). Given this observation, we concluded that pairwise differences in IBC are not due to differences in the duration of social interaction. Accordingly, group differences in time spent interacting are unlikely to fully explain group-level differences in IBC.

Since variation in IBC within a session could, in principle, obscure variation between sessions due to the dynamic and relationship-specific nature of social interaction, we next evaluated how IBC and behavior evolve throughout a given session. We computed IBC and the average inter-animal distance in 5-minute bins and found that both varied throughout a single recording session. We performed a mixed ANOVA for IBC by time bin (RM) and relationship type (naïve, partner, stranger). We observed a main effect of time, with IBC strongest at the start of a session (Fig. 2E; main effect of time F_3,123_ = 18.98, p-GG-corr =6.2e-9; Holm post-hoc: Bins 1 vs 2: T = 5.64, p-corr = 7e-6; Bins 1 vs 3: T = 5.05, p-corr = 3.5e-5; Bins 1 vs 4: T = 5.41, p-corr = 1.3e-5). We also replicated the effect of relationship on IBC (main effect of relationship: F_2,40_ = 3.54, p = 0.038).

Naïve and partners showed stronger IBC than strangers (Tukey HSD: Naïve vs Strangers p-adj = 0.001, Partners vs Strangers: p-adj = 0.014). We next examined social proximity (inter-vole distance) by time bin and relationship type. The average distance between animals varied significantly across time bins (Fig. 2F; F_3,120_ = 5.1312, p = 0.0106), and we observed a significant main effect of relationship type (F_2,40_ = 3.21, p = 0.026), such that partners were closer to each other, on average, than other types of dyads (Tukey’s HSD: naive-partners: mean-diff = −9.69, p < 0.0001; strangers-partners: mean-diff = −8.12, p = 0.0004).

### Bonding drives divergence in predictors of IBC between social contexts

Given that our analysis of single variables was insufficient to explain variation in IBC, we next used a linear mixed model (LMM) to identify the relational and real-time factors that contribute to IBC. We used IBC (Pearson’s r) as the dependent variable, calculated throughout the session with a 120-sec sliding window in 30-sec steps. The median distance between animals over the same sliding window (inter-animal distance), the average partner preference score of the interacting animals (bond strength), and time since the start of the interaction session/divider removal (time) were used as continuous predictors for IBC. To aid between-variable comparisons and intercept interpretation, all continuous variables were Z-scored, and time was centered at the session onset. Therefore, all model estimates reflect the normalized effects of each predictor at session onset (0 seconds) when animals are at the mean inter-animal distance (17.15 cm) and have a mean partner preference score (81.02%, experimental pairwise average) unless otherwise specified. To account for repeated measures, we included a random effect of recording session (N = 53 sessions, 2,286 observations). Relationship type was included as a categorical predictor to determine how these continuous predictors varied based on the relationship between animals. Log-ratio tests supported the exclusion of the 4th order interaction term and the inter-animal distance x bond strength x relationship type term. All other terms were retained, as they significantly improved the model’s performance (p < 0.05) (see Methods for additional details).

Our primary goal was to determine how social bonding alters the relationship between IBC and its behavioral predictors. To isolate bonding-related effects, we initially included all four relationship groups and compared strangers and partners after pairing to the naïve 1 and naïve 2 (future partner) groups. We expected minimal differences between groups before pairing, allowing this comparison to serve as a pseudo-control for the paired groups. We anticipated that the only differences in these two naïve groups would be in the effect of bond strength—specifically, IBC with a future partner before bonding could potentially predict bond strength after pairing. However, we did not see significant differences in the effects of bond strength (or any other predictors) on IBC between the naïve groups (Supp. Fig. 2D-G, Table 1; β = −0.005, SE = 0.07, z = −0.08, p = 0.934, all other p > 0.36). The highly similar effects of all predictors in these groups confirm that any post-pairing differences are unlikely to reflect pre-existing group differences. Given the consistent findings in the naïve group, we reran the model with these groups combined into a single naïve group to allow direct comparisons between IBC before and after pairing. Both models performed significantly above chance (Supp. Fig. 2B and C; Permutation tests: Four-group model, Log-Likelihood: −261.93, p < 0.0001; combined naïve group model, Log-Likelihood: −262.76, p < 0.0001).

**Table 1.** Linear mixed-effects model relating interbrain correlation (IBC) to distance, time, bond strength, and relationship type with comparisons between naive groups. The table reports model coefficients (Coef.), standard errors (SE), z statistics, two-sided *P* values (P>|z|), and 95% confidence intervals for fixed effects. The naïve future partners group served as the reference level for relationship type. Relationship interaction terms represent differences in slopes relative to this reference group. Distance (inter-animal distance), time (time since session onset), and bond strength (average partner preference score of the dyad) were normalized prior to analysis. Variables were centered so that terms reflect values at session onset, at mean distance (17.15cm), and mean bond strength (81.02%). A random intercept was included for recording (variance shown).

|  | Coef. | SE | z | P>\|z\| | [0.025 | 0.975] |
| --- | --- | --- | --- | --- | --- | --- |
| Intercept | 0.354 | 0.048 | 7.38 | 0 | 0.26 | 0.448 |
| Relationship[T.Naive 1] | -0.06 | 0.066 | -0.911 | 0.362 | -0.19 | 0.069 |
| Relationship[T.Partners] | 0.1 | 0.074 | 1.343 | 0.179 | -0.046 | 0.246 |
| Relationship[T.Strangers] | -0.194 | 0.07 | -2.794 | 0.005 | -0.331 | -0.058 |
| Distance | -0.054 | 0.032 | -1.696 | 0.09 | -0.117 | 0.008 |
| Relationship[T.Naive 1]:Distance | 0.02 | 0.041 | 0.472 | 0.637 | -0.062 | 0.101 |
| Relationship[T.Partners]:Distance | 0.229 | 0.063 | 3.657 | 0 | 0.106 | 0.352 |
| Relationship[T.Strangers]:Distance | -0.1 | 0.052 | -1.926 | 0.054 | -0.202 | 0.002 |
| Time | -0.045 | 0.012 | -3.694 | 0 | -0.069 | -0.021 |
| Relationship[T.Naive 1]:Time | 0.004 | 0.017 | 0.239 | 0.811 | -0.029 | 0.037 |
| Relationship[T.Partners]:Time | -0.086 | 0.024 | -3.554 | 0 | -0.133 | -0.038 |
| Relationship[T.Strangers]:Time | 0.035 | 0.019 | 1.869 | 0.062 | -0.002 | 0.072 |
| Bond Strength | 0.076 | 0.052 | 1.456 | 0.145 | -0.026 | 0.179 |
| Bond Strength:Relationship[T.Naive 1] | -0.005 | 0.066 | -0.082 | 0.934 | -0.135 | 0.124 |
| Bond Strength:Relationship[T.Partners] | 0.13 | 0.069 | 1.884 | 0.059 | -0.005 | 0.266 |
| Bond Strength:Relationship[T.Strangers] | -0.134 | 0.078 | -1.724 | 0.085 | -0.287 | 0.018 |
| Bond Strength:Time | 0.002 | 0.013 | 0.142 | 0.887 | -0.024 | 0.028 |
| Bond Strength:Relationship[T.Naive 1]:Time | -0.009 | 0.016 | -0.546 | 0.585 | -0.041 | 0.023 |
| Bond Strength:Relationship[T.Partners]:Time | -0.065 | 0.019 | -3.513 | 0 | -0.101 | -0.029 |
| Bond Strength:Relationship[T.Strangers]:Time | 0.057 | 0.02 | 2.833 | 0.005 | 0.018 | 0.096 |
| Distance:Time | 0.023 | 0.015 | 1.552 | 0.121 | -0.006 | 0.052 |
| Relationship[T.Naive 1]:Distance:Time | -0.004 | 0.019 | -0.212 | 0.832 | -0.041 | 0.033 |
| Relationship[T.Partners]:Distance:Time | -0.102 | 0.029 | -3.533 | 0 | -0.159 | -0.045 |
| Relationship[T.Strangers]:Distance:Time | 0.001 | 0.021 | 0.039 | 0.969 | -0.041 | 0.042 |
| Bond Strength:Distance | 0.038 | 0.018 | 2.107 | 0.035 | 0.003 | 0.073 |
| Bond Strength:Distance:Time | -0.019 | 0.008 | -2.378 | 0.017 | -0.034 | -0.003 |
| Recording Variance | 0.022 | 0.018 |  |  |  |  |

Overall, the model predicts differences in the link between IBC and the predictors that depend on the relationship between the interacting voles (Fig. 2G-I). Accounting for the interactions between these predictors elucidated relationships between IBC, bond strength, and inter-animal distance that were not captured by single-variable analyses. Specifically, we found that pairing led to a divergence in IBC and its relationship to these predictors for partner and stranger interactions, compared to interactions between sexually naïve individuals (Fig. 2G-I). Having accounted for both bond strength and inter-animal distance, we found that each group still had positive baseline IBC, supporting our finding that IBC is a conserved feature of social interaction broadly (Table 2; Naïve: β = 0.324, Strangers: β = 0.159, Partners: β = 0.452). Unlike our prior comparisons of IBC across the entire recording session, this model revealed that pairing led to a significant increase in IBC between partners, compared to the naïve group, and a decrease in IBC between strangers (Table 2; naïve vs partners: β = 0.128, SE = 0.07, z = 1.98, p < 0.048, naïve vs strangers: β = −0.165, SE = 0.06, z = −2.73, p = 0.006).

**Table 2.** Linear mixed-effects model relating interbrain correlation (IBC) to distance, time, and bond strength with naïve groups combined. The table reports model coefficients (Coef.), standard errors (SE), z statistics, two-sided p-values (P>|z|), and 95% confidence intervals for fixed effects. The combined naïve group served as the reference level for relationship type. Relationship interaction terms represent differences in slopes relative to this reference group. Distance (inter-animal distance), time (time since session onset), and bond strength (average partner preference score of the dyad) were normalized prior to analysis. Variables were centered so that terms reflect values at session onset, at mean distance (17.15cm), and mean bond strength (81.02%). A random intercept was included for recording (variance shown).

|  | Coef. | SE | z | P>\|z\| | [0.025 | 0.975] |
| --- | --- | --- | --- | --- | --- | --- |
| Intercept | 0.324 | 0.033 | 9.866 | 0 | 0.26 | 0.389 |
| Relationship [T.Partners] | 0.128 | 0.065 | 1.976 | 0.048 | 0.001 | 0.256 |
| Relationship[T.Strangers] | -0.165 | 0.06 | -2.728 | 0.006 | -0.283 | -0.046 |
| Distance | -0.044 | 0.019 | -2.306 | 0.021 | -0.081 | -0.007 |
| Relationship[T.Partners]:Distance | 0.218 | 0.055 | 3.962 | 0 | 0.11 | 0.325 |
| Relationship[T.Strangers]:Distance | -0.11 | 0.045 | -2.444 | 0.015 | -0.199 | -0.022 |
| Time | -0.043 | 0.008 | -5.122 | 0 | -0.059 | -0.026 |
| Relationship[T.Partners]:Time | -0.088 | 0.022 | -3.957 | 0 | -0.131 | -0.044 |
| Relationship[T.Strangers]:Time | 0.033 | 0.016 | 1.993 | 0.046 | 0.001 | 0.065 |
| Bond Strength | 0.075 | 0.032 | 2.333 | 0.02 | 0.012 | 0.138 |
| Bond Strength:Relationship[T.Partners] | 0.132 | 0.056 | 2.345 | 0.019 | 0.022 | 0.241 |
| Bond Strength:Relationship[T.Strangers] | -0.133 | 0.066 | -2.006 | 0.045 | -0.263 | -0.003 |
| Bond Strength:Time | -0.004 | 0.008 | -0.447 | 0.655 | -0.019 | 0.012 |
| Bond Strength:Relationship[T.Partners]:Time | -0.059 | 0.016 | -3.797 | 0 | -0.09 | -0.029 |
| Bond Strength:Relationship[T.Strangers]:Time | 0.063 | 0.017 | 3.641 | 0 | 0.029 | 0.096 |
| Distance:Time | 0.021 | 0.009 | 2.472 | 0.013 | 0.004 | 0.038 |
| Relationship[T.Partners]:Distance:Time | -0.099 | 0.026 | -3.846 | 0 | -0.15 | -0.049 |
| Relationship[T.Strangers]:Distance:Time | 0.003 | 0.017 | 0.155 | 0.876 | -0.031 | 0.037 |
| Bond Strength:Distance | 0.036 | 0.017 | 2.151 | 0.031 | 0.003 | 0.07 |
| Bond Strength:Distance:Time | -0.019 | 0.007 | -2.471 | 0.013 | -0.033 | -0.004 |
| Recording Variance | 0.022 | 0.018 |  |  |  |  |

We also found that correlations between IBC and bond strength depended on the relationship type. First, stronger IBC during naïve interactions predicted bond strength after pairing (*i.e.,* a stronger average partner preference; Fig. 2G, Table 2; β = 0.075, SE = 0.03, z = 2.33, p < 0.02). This finding suggests that voles with stronger IBC during initial opposite-sex introductions (with any potential future mate) have a stronger propensity to form pair bonds, likely reflecting generalized prosocial and affiliative tendencies. After bonding, we observed an even stronger positive correlation between IBC and bond strength with partners, compared to naïve individuals, and a modest negative correlation in strangers (Fig. 2G, Table 2; naïve vs partners: β = 0.132, SE = 0.06, z = 2.35, p = 0.019, naïve vs strangers: β = −0.133, SE = 0.07, z = −2.01, p = 0.045). Thus, after pairing, the selectivity of the bond is reflected in IBC during social interactions. This contrasts with the non-selective, general prosocial signal in IBC before pairing that is predictive of later bond strength.

Our model further revealed that, before pairing, IBC decreased with inter-animal distance, which provides a coarse proxy for social interaction (Fig. 2H; Table 2; β = −0.044, SE = 0.02, z = −2.31, p = 0.021). This effect was even stronger during stranger interactions after pairing (naïve vs strangers: β = −0.11, SE = 0.05, z = −2.44, p = 0.015). These findings suggest that IBC is stronger when animals are close enough to interact, consistent with prior findings in mice and bats. However, this finding did not hold for the partners group, which showed the opposite trend; greater inter-animal distances predicted stronger IBC (Fig. 2H; naïve vs partners: β = 0.218, SE = 0.06, z = 3.96, p > 0.001). While unexpected, this may be explained by behavioral differences between groups, with partners spending most of their time in very close proximity, reducing the efficacy of proximity as a measure of active social interaction (Fig. 2F). Bonded voles are known to spend a significant portion of their time huddling^36^. During huddling, voles are in very close proximity but often resting, rather than actively engaging with one another, which may explain the negative relationship between IBC and proximity. Regardless, our data indicate that IBC dynamics differ when interacting with a bonded partner versus an unknown individual, suggesting that bond formation leads to modulation of the factors that predict IBC.

Finally, IBC decreased throughout the course of the session for all groups, likely due to the high initial salience of introduction and reintroduction. This salience decreases, along with active social investigation, as the animals habituate to one another (Fig. 2I; Table 2; Naïve: β = −0.043, SE = 0.01, z = −5.12, p < 0.001). The effect was pronounced in partners and relatively weak for strangers (naïve vs partners: β = −0.088, SE = 0.02, z = −3.96, p < 0.001; naïve vs strangers: β = 0.033, SE = 0.02, z = 1.99, p = 0.046). This may reflect increased salience of partner reunion and/or a more rapid habituation, given their familiarity. Time also attenuated all relationship-specific effects of bond strength and distance on IBC (Fig 2. J-L; Table 2). For example, the negative association between inter-animal distance and IBC seen in naïve and stranger dyads became less negative over time; likewise, the positive relationship between bond strength and IBC in partners and naïve animals attenuated over time. Thus, like IBC, the effects of bond strength and distance on IBC also depend on active and salient social interaction.

### Machine learning-based behavioral classification in freely interacting voles

While inter-animal distance significantly predicted IBC in the LMM, proximity alone represents an extremely coarse proxy for social interaction, failing to capture the diversity of social behavior expressed between two animals. For instance, two animals can be close to each other but exhibit aggression or affiliation. To obtain refined and less biased behavioral annotation, we developed a machine learning-based pipeline to classify social behaviors in freely interacting prairie voles.

Our behavior classification pipeline integrates pose estimation, feature extraction, and hierarchical clustering across 86 videos of same-sex and opposite-sex pairs spanning multiple relationship types, ultimately identifying 35 discrete social behaviors (Fig. 3, Supp. Fig. 3). We first applied DeepLabCut to track anatomically defined body landmarks for both animals^37^ (Fig. 3B; see Methods for details). From these tracked coordinates, we extracted a set of features designed to capture inter-animal configurations and their dynamics (Fig. 3C-E). To enable scalable clustering across ∼1.6 million frames, we implemented a two-stage dimensionality reduction and clustering strategy. First, a high-efficiency algorithm grouped frames with similar feature vectors, reducing the dataset to ∼100,000 representative clusters^38^. Second, Ward agglomerative clustering was applied to the reduced feature space, generating a hierarchical dendrogram (Fig. 3I, Supp. Fig. 3a)^39^.

**Figure 3.**
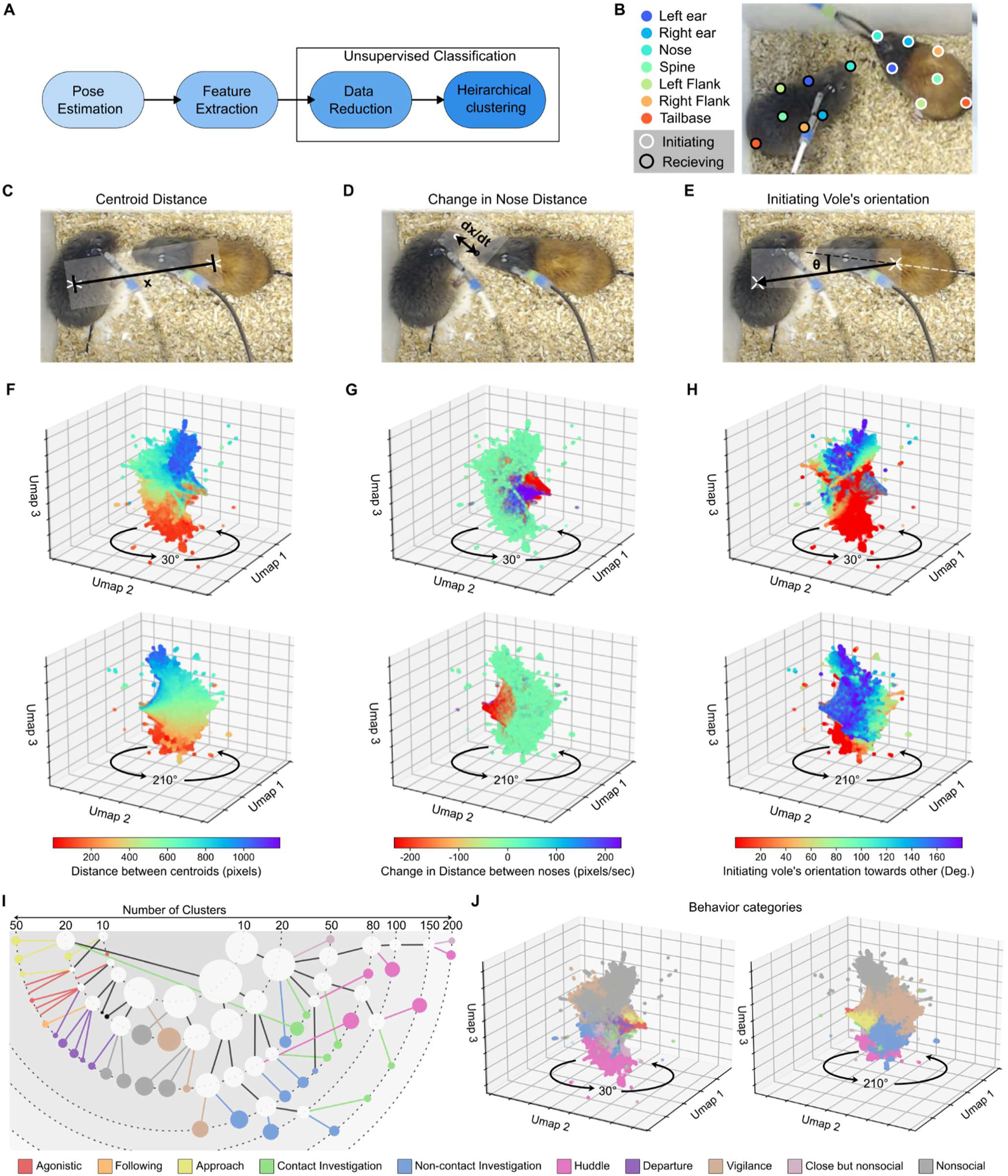
Machine-learning based classification of social behavior in prairie vole dyads. **A.** Behavior classification process. **B.** Example key-point labeling used in multi-animal DeepLabCut. **C-E.** Examples of calculated features. **F-H.** UMAP visualizations showing the feature value for each frame included in clustering from two different angles. Features show distinct distributions of values across the UMAP. **I.** The hierarchical clustering process created a tree that was examined at multiple points based on the number of clusters. Cluster color corresponds to behavioral category; white clusters were further differentiated by identifying subclusters that contained only one behavior. **J.** UMAP visualization color coded for the behavior identified in each frame. Behavior categories correspond to expected features for those frames when compared with F-H.

Behavioral categories were defined by identifying cut heights within the dendrogram where clusters formed behaviorally coherent and clearly differentiated groupings. This approach yielded 35 distinct motifs, which were sorted into 10 behavior categories. Behavioral categories included contact investigation and non-contact investigation, differentiated in motifs based on relative orientation and, for the non-contact investigations, distance as well (nose-to-nose or nose-to-tail). The clustering also revealed motifs showing agonistic behaviors (*e.g*., lunging, chasing), social approach, and social vigilance, where one individual monitors the other from a distance^40^ (Fig. 3I, Supp. Fig. 3a). We assessed the correspondence between assigned behavioral categories and their defining features by embedding frames into UMAP space and overlaying feature values and cluster assignments (Fig. 3F-H, and J, Supp. Fig. 3B). This visualization revealed a clear structure consistent with behavioral intuitions. For example, inter-animal distance was lowest in clusters corresponding to huddling and highest in non-social clusters (Fig. 3F and J). Similarly, approach and departure mapped cleanly to frames with negative and positive changes in inter-nose distances, respectively (Fig. 3G and J).

### Behaviors differ by relationship type

We annotated behavior in a subset of interbrain recordings optimized for our behavior classification method (see Methods for details, n = 32 recordings from 8 pairs). Using the 10 behavioral classes (Fig. 3J), we first asked whether each relationship (naïve, partner, and stranger) was associated with different behaviors during free interaction. As expected, the behavioral composition of interactions varied across the three relationship groups (PERMANOVA, pseudo-F_2, 29_ 2.77, p = 0.017, 999 permutations) (Fig. 4A). Post hoc tests confirmed that partners behaved significantly different than naive animals or strangers and that naïve and stranger dyads where not significantly different from each other (PERMANOVA, 9,999 permutations, Bonferroni-corrected: partners vs naïve: pseudo-F_1, 22_ = 4.08, p-adj = 0.029; partners vs strangers: pseudo-F_1,14 =_ 3.85, p-adj = 0.036; naïve vs strangers: pseudo-F_1,22_ = 0.65, p-adj > 1). The results were the same when all 35 behavioral motifs instead of the broader behavioral categories (main-effect: pseudo-F_2,29_ = 3.17, p = 0.008; partners vs naive: pseudo-F_1,22_ = 5.06, p-adj = 0.015; partners vs strangers: pseudo-F_1,14_ = 3.96, p-adj = 0.018; naive vs strangers: pseudo-F_1,22_ = 0.76, p-adj > 1). Based on visual inspection, group-level differences in behavioral composition were driven, at least in part, by partners spending much more time huddling compared to the other two groups (Fig. 4A), consistent with past work in this species^33,41,42^.

**Figure 4.**
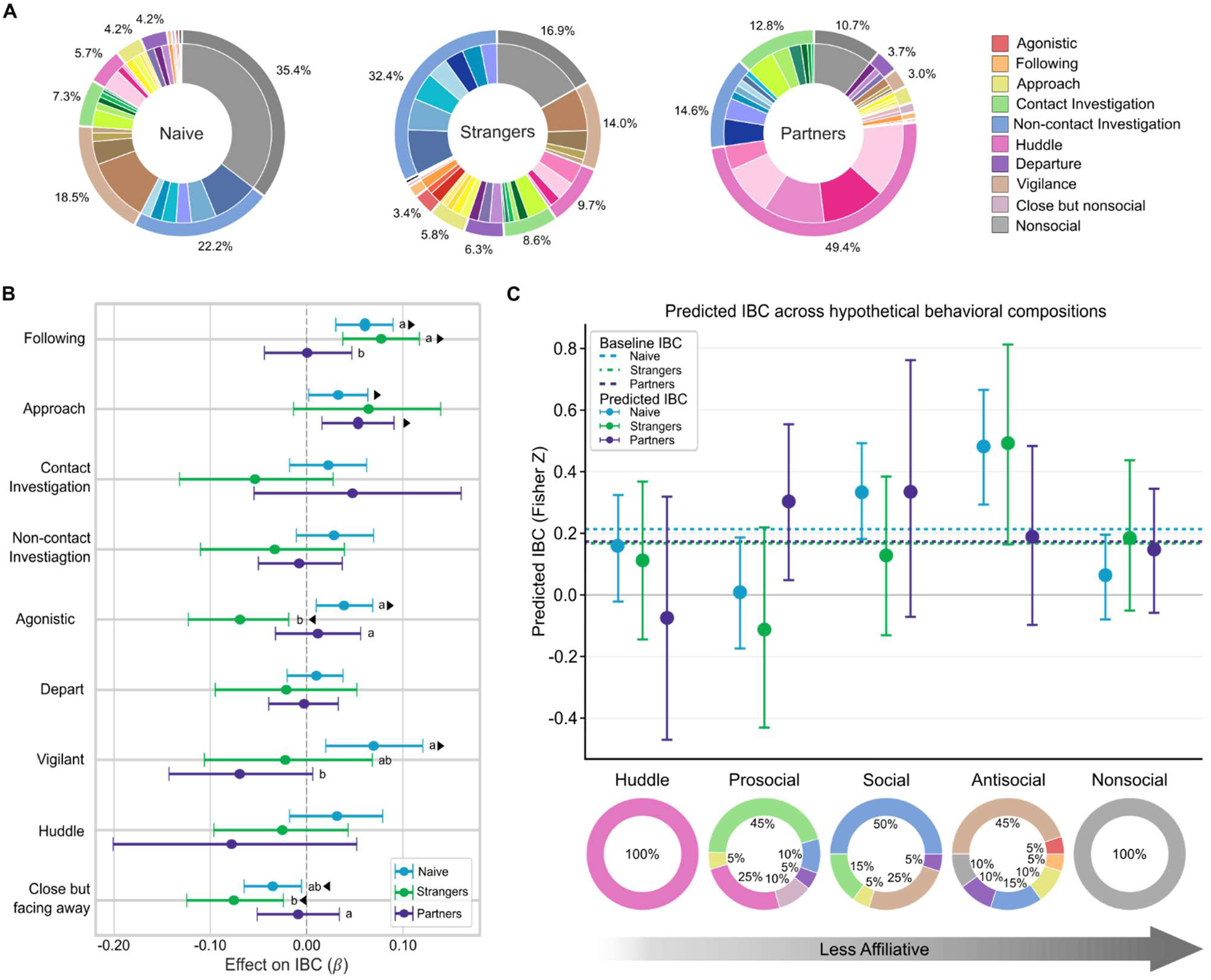
Behavior and IBC associations were modulated by the relationship type. **A**. Naïve, stranger, and partner dyads differ in the distribution of behaviors (PERMANOVA: effect of relationship: p = 0.017). Partners displayed significantly different behavior than naïve voles or strangers (Bonferroni-corrected: Partners vs Naïve: p-adj = 0.029, partners vs strangers: p-adj = 0.036). **B.** Forest plot showing *β* values for the relationship between each normalized behavior Center Log Ratio value and IBC. Error bars show the 95% HDI intervals compared to zero. Right and left pointed triangles denote *β* values that are credibly different than zero. Letters denote credible differences between relationship type *β* values for each specific behavior. **C.** Predicted IBC values for each relationship type for different 120-sec behavior compositions. Pie charts along the x-axis show the hypothetical behavior composition used to generate the predicted IBC value, from most to least affiliative. Horizontal lines show predicted baseline IBC (Model Intercepts for each group).

### Relationship type modulates the association between IBC and behaviors

To quantify how behavior predicted IBC, we fit a Bayesian hierarchical regression model. We used the same sliding window IBC calculation as in the above LMMs. We included the proportion of time spent in each behavior category across the sliding window, the relationship between individuals (naive, stranger, partner), and the interactions between the relationship and each behavior as predictors. We used a centered log ratio transform on behavioral proportions to account for their compositional structure, then standardized prior to modeling. To avoid perfect collinearity, we excluded non-social behavior from our predictors, having it serve as the default behavioral category. A Bayesian framework allowed partial pooling across recordings and stabilized parameter estimates, which was important given the variability in behavioral distributions across relationship groups. We did not include time or bond strength in this model because their higher-order interactions with relationship type and behavior would have resulted in substantial parameter expansion relative to the available sample size. The present model was therefore specified to isolate relationship type-dependent effects of behavioral composition on IBC. To validate the model, we fitted models on permuted data (500 permutations; see Methods for details); comparisons of log likelihood and R^2^ values showed that our model performed above chance (Supp. Fig. 4c; permutation test: p = 0.002).

The Bayesian model revealed that the effect of behavior on IBC was modulated by the relationship between individuals (Fig. 4B, Table 3). First, we determined which behaviors affected IBC for each relationship group using posterior contrasts. These effects were considered credible when the 95% highest density interval (HDI) excluded zero. For example, ‘approach’ was positively associated with IBC in naïve and partner dyads and exhibited a similar effect in strangers, suggesting that ‘approach’ broadly corresponds to enhanced IBC (naïve: mean = 0.033, 95% HDI [0.002, 0.064]; partners: mean = 0.054, 95% HDI [0.017, 0.092]; strangers: mean = 0.064, 95% HDI [−0.015, 0.135]). We also used posterior contrasts to ask how the effects of behavior on IBC differed between relationship types. For example, ‘following’ had a credibly stronger effect on IBC in naive and stranger dyads than in partners (naïve vs partners: mean = −0.06, 95% HDI [−0.118, −0.009]; strangers vs partners: mean = −0.076, 95% HDI [−0.139, −0.017]), while ‘agonistic’ behavior was positively correlated with IBC in naïve dyads but negatively correlated with IBC in stranger dyads, with little effect in partners (naïve: mean = 0.039, 95% HDI [0.010, 0.069]; strangers: mean = - 0.070, 95% HDI [−0.122, −0.020], partners: mean = 0.012, 95% HDI [−0.032, 0.056]). Further, IBC between strangers during agonistic behavior was credibly lower than in naïve and partner dyads (strangers vs naïve: mean = −0.109, 95% HDI [−0.166, −0.049]; strangers vs partners: mean = - 0.082, 95% HDI [−0.148, −0.010]). These group-level differences in behavior and IBC associations demonstrate that relationship-dependent differences in IBC do not simply reflect relationship-dependent differences in behavioral expression.

**Table 3.** Posterior estimates from the Bayesian model relating behavioral state and relationship type to interbrain correlation (IBC). The table reports posterior means, standard deviations (SD), and 94% highest density intervals (HDI; 3–97%) for all model parameters. Fixed effects include CLR-transformed proportions for each behavioral category and relationship type, as well as their interactions. Naïve animals served as the reference group for relationship type. Random intercepts were included for each recording. Diagnostic statistics are also shown, including the Monte Carlo standard error of the mean (MCSE), effective sample sizes (ESS) for the bulk and tail of the posterior distributions, and the potential scale reduction factor (R^). All parameters showed good convergence (R^ ≈ 1).

| | mean | sd | HDI<br>3% | HDI<br>97% | MCSE<br>mean | MCSE<br>sd | ESS<br>bulk | ESS<br>tail | $\hat{R}$ |
| --- | --- | --- | --- | --- | --- | --- | --- | --- | --- |
| sigma | 0.207 | 0.006 | 0.196 | 0.218 | 0 | 0 | 11204 | 5329 | 1 |
| Intercept | 0.202 | 0.063 | 0.087 | 0.324 | 0.001 | 0.001 | 2763 | 4059 | 1 |
| Relationship[Strangers] | -0.03 | 0.114 | -0.262 | 0.171 | 0.002 | 0.001 | 3807 | 4847 | 1 |
| Relationship[Partners] | -0.035 | 0.108 | -0.24 | 0.167 | 0.002 | 0.001 | 3453 | 4511 | 1 |
| Vigilant | 0.048 | 0.029 | -0.007 | 0.102 | 0 | 0 | 6223 | 6356 | 1 |
| Approach | 0.03 | 0.017 | -0.002 | 0.061 | 0 | 0 | 7910 | 6504 | 1 |
| Noncontact investigation | 0.032 | 0.024 | -0.01 | 0.079 | 0 | 0 | 6285 | 5641 | 1 |
| depart | 0.017 | 0.016 | -0.013 | 0.045 | 0 | 0 | 8221 | 6445 | 1 |
| Contact investigation | 0.002 | 0.023 | -0.041 | 0.044 | 0 | 0 | 6619 | 6389 | 1 |
| Following | 0.04 | 0.017 | 0.008 | 0.073 | 0 | 0 | 7573 | 6276 | 1 |
| Agonistic | 0.044 | 0.016 | 0.012 | 0.073 | 0 | 0 | 8002 | 6302 | 1 |
| Huddle | 0.022 | 0.027 | -0.029 | 0.072 | 0 | 0 | 9142 | 6663 | 1 |
| Close facing away | -0.036 | 0.017 | -0.068 | -0.004 | 0 | 0 | 6359 | 6038 | 1 |
| Rel:Vigilant[Strangers] | -0.045 | 0.054 | -0.147 | 0.058 | 0.001 | 0 | 7659 | 6480 | 1 |
| Rel:Vigilant[Partners] | -0.118 | 0.052 | -0.213 | -0.018 | 0.001 | 0 | 6765 | 6409 | 1 |
| Rel:Approach[Strangers] | 0.053 | 0.043 | -0.023 | 0.137 | 0 | 0 | 7517 | 6579 | 1 |
| Rel:Approach[Partners] | 0.054 | 0.029 | -0.002 | 0.106 | 0 | 0 | 9087 | 6651 | 1 |
| Rel:Noncontact investigation [Strangers] | -0.061 | 0.042 | -0.138 | 0.018 | 0.001 | 0 | 6056 | 6093 | 1 |
| Rel:Noncontact investigation [Partners] | -0.043 | 0.033 | -0.105 | 0.018 | 0 | 0 | 7147 | 6275 | 1 |
| Rel:Depart[Strangers] | -0.048 | 0.04 | -0.124 | 0.023 | 0 | 0 | 8786 | 6913 | 1 |
| Rel:Depart[Partners] | -0.027 | 0.026 | -0.078 | 0.02 | 0 | 0 | 8535 | 6250 | 1 |
| Rel:Contact investigation[Strangers] | -0.046 | 0.046 | -0.136 | 0.035 | 0.001 | 0 | 7133 | 6055 | 1 |
| Rel:Contact investigation[Partners] | 0.048 | 0.066 | -0.077 | 0.171 | 0.001 | 0.001 | 8272 | 6592 | 1 |
| Rel:Following[Strangers] | 0.012 | 0.029 | -0.042 | 0.067 | 0 | 0 | 8542 | 6303 | 1 |
| Rel:Following[Partners] | -0.033 | 0.03 | -0.087 | 0.024 | 0 | 0 | 9215 | 6638 | 1 |
| Rel:Agonistic[Novels] | -0.096 | 0.034 | -0.157 | -0.03 | 0 | 0 | 9036 | 6877 | 1 |
| Rel:Agonistic[Strangers] | -0.033 | 0.029 | -0.088 | 0.021 | 0 | 0 | 8168 | 6800 | 1 |
| Rel:Huddle[Strangers] | -0.031 | 0.049 | -0.121 | 0.063 | 0 | 0 | 9878 | 6647 | 1 |
| Rel:Huddle[Partners] | -0.096 | 0.078 | -0.242 | 0.051 | 0.001 | 0.001 | 7336 | 5715 | 1 |
| Rel:Close facing away [Strangers] | -0.028 | 0.032 | -0.089 | 0.031 | 0 | 0 | 7963 | 6743 | 1 |
| Rel:Close facing away [Partners] | 0.042 | 0.03 | -0.015 | 0.098 | 0 | 0 | 7827 | 6527 | 1 |
| 1\|Recording_sigma | 0.204 | 0.037 | 0.139 | 0.275 | 0.001 | 0 | 3153 | 4676 | 1 |
| 1\|Recording[1] | 0.012 | 0.075 | -0.123 | 0.157 | 0.001 | 0.001 | 3499 | 4726 | 1 |
| 1\|Recording[2] | 0.08 | 0.108 | -0.12 | 0.284 | 0.001 | 0.001 | 5505 | 5634 | 1 |
| 1\|Recording[3] | -0.182 | 0.073 | -0.318 | -0.044 | 0.001 | 0.001 | 3469 | 4854 | 1 |
| 1\|Recording[4] | 0.299 | 0.094 | 0.121 | 0.471 | 0.001 | 0.001 | 4452 | 5471 | 1 |
| 1\|Recording[5] | 0.274 | 0.103 | 0.08 | 0.464 | 0.001 | 0.001 | 5142 | 5565 | 1 |
| 1\|Recording[6] | 0.147 | 0.073 | 0.005 | 0.279 | 0.001 | 0.001 | 3340 | 5098 | 1 |
| 1\|Recording[7] | 0.109 | 0.072 | -0.018 | 0.254 | 0.001 | 0.001 | 3352 | 4427 | 1 |
| 1\|Recording[8] | 0.17 | 0.094 | -0.004 | 0.349 | 0.001 | 0.001 | 4557 | 5667 | 1 |
| 1\|Recording[9] | -0.161 | 0.105 | -0.367 | 0.027 | 0.001 | 0.001 | 5657 | 5515 | 1 |
| 1\|Recording[10] | 0.036 | 0.089 | -0.133 | 0.201 | 0.001 | 0.001 | 4055 | 4964 | 1 |
| 1\|Recording[11] | -0.131 | 0.078 | -0.279 | 0.013 | 0.001 | 0.001 | 3585 | 5153 | 1 |
| 1\|Recording[12] | 0.033 | 0.071 | -0.1 | 0.167 | 0.001 | 0.001 | 3333 | 4655 | 1 |
| 1\|Recording[13] | -0.057 | 0.091 | -0.222 | 0.119 | 0.001 | 0.001 | 4291 | 5108 | 1 |
| 1\|Recording[14] | -0.179 | 0.073 | -0.325 | -0.049 | 0.001 | 0.001 | 3478 | 4852 | 1 |
| 1\|Recording[15] | -0.183 | 0.102 | -0.375 | 0.006 | 0.001 | 0.001 | 5423 | 5212 | 1 |
| 1\|Recording[16] | -0.056 | 0.073 | -0.198 | 0.075 | 0.001 | 0.001 | 3433 | 5276 | 1 |
| 1\|Recording[17] | -0.251 | 0.091 | -0.415 | -0.071 | 0.001 | 0.001 | 4085 | 4767 | 1 |
| 1\|Recording[18] | -0.131 | 0.072 | -0.27 | 0.003 | 0.001 | 0.001 | 3404 | 5217 | 1 |
| 1\|Recording[19] | 0.007 | 0.104 | -0.187 | 0.204 | 0.001 | 0.001 | 5335 | 5440 | 1 |
| 1\|Recording[20] | 0.05 | 0.073 | -0.081 | 0.192 | 0.001 | 0.001 | 3234 | 4783 | 1 |
| 1\|Recording[21] | 0.386 | 0.074 | 0.247 | 0.525 | 0.001 | 0.001 | 3469 | 4753 | 1 |
| 1\|Recording[22] | -0.189 | 0.092 | -0.361 | -0.016 | 0.001 | 0.001 | 4277 | 5516 | 1 |

**Table 4.** Behavioral classification feature set. The names and definitions for the feature set used in the unsupervised classification. Per-vole metrics were calculated for both the initiating and receiving vole, resulting in two features per measure. Each feature was given a weight for clustering, depending on its importance in distinguishing behaviors.

| Name | Definition | Type | Weight |
| --- | --- | --- | --- |
| Distance Between Voles | Euclidean distance between median centroids. | Inter-vole | 2 |
| Change in Distance | Frame-to-frame change in centroid distance (positive = moving apart, negative = moving closer). | Inter-vole | 1 |
| Nose-to-Nose Distance | Distance between each vole's nose | Inter-vole | 1 |
| Change in Nose Distance | Frame-to-frame change in centroid distance (positive = moving apart, negative = moving closer). | Inter-vole | 0.5 |
| Speed | Median Euclidean displacement of each body part between frames. | Per vole | 1 |
| Speed Variance | The variance of the above metric over a 10-frame window. | Per vole | 0.5 |
| Orientation Towards the Other | Angle between a fitted body vector and the line between the centroids of each vole (angles < 90° = facing towards, angles > 90° = facing away). | Per vole | 1 |
| Change in Orientation | Frame-to-frame change in the global orientation smoothed over a 10-frame window | Per vole | 0.5 |
| Speed Towards the Other | Projection of the median velocity vector onto the line between the centroids of each vole. | Per vole | 0.75 |

To further examine the role of relationship in modulating IBC, we generated predictions of IBC for each relationship type under representative 120-sec behavioral compositions that spanned a range of affiliative to antisocial and non-social behavior: huddling, prosocial, social, antisocial, and nonsocial (Fig. 4C). These reflected commonly observed compositions of behavior across all relationship types in our dataset, with two notable exceptions: naïve pairs never huddled for 100% of any 120-sec window, and partners were never non-social for a full 120 seconds. Predictions were derived directly from the fitted model’s posterior distributions, allowing estimation of expected IBC and the associated uncertainty for each scenario. Baseline IBC for each group corresponded to the model intercept for each relationship group.

Huddling was associated with lower predicted IBC across all relationship groups, consistent with the passive nature of this behavior. In contrast, for the prosocial composition comprised primarily of contact investigation and huddling, the model predicted elevated IBC in partners relative to their baseline, whereas naïve pairs and strangers showed predicted values below baseline. Only naïve pairs showed an association between the moderately social composition (largely noncontact investigation with moderate vigilance) and elevated predicted IBC; strangers were not predicted to deviate from their baseline, while partners exhibited a wide range of predicted IBC values. Intriguingly, the antisocial behavior composition was associated with elevated predicted IBC in naive and stranger pairs, but values near baseline in partners, potentially consistent with these behaviors driving cognitive engagement in both members of the dyad, especially when the other individual is unknown. The nonsocial composition was associated with reduced predicted IBC in naïve pairs and values near baseline for partners and strangers.

## Discussion

A growing body of human hyperscanning research has established that IBC is a hallmark of social interaction, scaling with relationship quality, empathy, and cooperative behavior^18,43–46^. Yet, how IBC changes as social bonds form, and what neural and behavioral factors shape it, has remained largely inaccessible, given the difficulty of implementing controlled longitudinal designs in human attachment formation. Here, we demonstrate that monogamous prairie voles exhibit IBC in the medial prefrontal cortex during free social interaction, detectable via fiber photometry for GCaMP-mediated fluorescence. Prairie voles thus provide a unique and tractable model for interrogating how IBC evolves across the formation and maintenance of selective social bonds. Using a longitudinal design, we show that bond formation reshapes IBC in a relationship-specific manner, that individual variation in IBC is best explained by a combination of bond strength, inter-animal distance, and time since interaction onset, and that the relationship between social behavior and IBC is itself modulated by the nature of the relationship between individuals.

While romantic partners and close friends consistently show stronger IBC than strangers in human hyperscanning studies, those comparisons are cross-sectional and do not track IBC across the transition from stranger to partner within the same dyad. We also found that IBC with partners is stronger than with strangers when comparing values at the bonded time point. However, perhaps contrary to expectation, the overall difference in partner and stranger IBC is driven by decreased IBC between bonded animals and opposite-sex strangers, while IBC across the entire recording session is similar between partner dyads and pre-bonding interactions. Notably, we found that IBC with a bonded partner scales positively with bond strength after accounting for distance and time variables, corroborating the core finding from human studies that IBC indexes the strength of social attachment. Thus, pair bonding does not simply amplify IBC, but it reshapes it in a relationship-specific manner, strengthening the association between IBC and bond quality while simultaneously reducing synchrony with unfamiliar individuals.

IBC in the mPFC has previously been reported in mice and bats. In both species, the amount of social engagement during a session positively predicts IBC strength^28,29^. Contrary to expectations, we did not replicate this relationship for voles when looking at inter-animal distance in isolation. Even after accounting for bond-strength and time within the session, Inter-animal distance did not consistently predict IBC across all relationship types. In bonded partners, this relationship is reversed, with greater distance associated with stronger IBC at the session onset. This divergence may reflect a key ethological difference in baseline social engagement across species: voles in our study spent a median of approximately 85% of each session engaged in social interactions, compared to less than 50% in bats and roughly 15% in mice. When social interaction is nearly continuous, inter-animal distance becomes a poor proxy for the quality or type of social engagement. This motivated the development of a more granular approach to behavioral quantification.

To characterize and quantify diverse social behaviors, we developed one of the first machine-learning-based social behavior classification pipelines for freely interacting prairie voles (see also Bueno-Junior *et al*.^47^). We used the resulting behavioral annotation in subsequent Bayesian modeling to identify the association between different behaviors and IBC in a relationship-dependent manner. This approach revealed that the coupling between behavior and IBC was not fixed but depended on the relationship between animals. Remarkably, the exact same behavior can differentially modulate interbrain neural synchrony depending on the social interaction partner. One likely contributor to this social context dependence is the differential salience of interactions with known and unknown individuals.

Bonding represents a fundamental state change: voles transition from a state where all unknown opposite-sex individuals represent potential partners to one where such individuals are actively rejected. This means that initial interactions with unknown opposite-sex voles are highly salient both before and after bonding, but for fundamentally different reasons, reflected in different behaviors. Supporting this idea, subordinate mice show the greatest number of neurons tracking the behavior of dominant individuals, suggesting that heightened social vigilance is directly reflected in interbrain neural synchrony^28^. Differential salience of the same behaviors, driven by the nature of the relationship between animals, could therefore account for the differences in behavioral modulation of IBC. To illustrate such relationship-dependent effects concretely, we simulated IBC predictions across hypothetical behavioral scenarios commonly observed in our recordings. Prosocial interaction was associated with the greatest IBC in bonded partners, while naïve and stranger pairs showed elevated IBC during moderately social and antisocial scenarios, respectively. This overall pattern suggests that bond formation does not merely change animals’ behaviors, but that differences in social context fundamentally change the connection between behavior and emergent neural synchrony.

We propose the following framework based on our results. For bonded partners, shared history may enable a more accurate prediction of a partner’s behavior and internal state. This, coupled with coordinated behaviors to achieve shared goals, enhances IBC. We expect this to be most evident during prosocial, affiliative interactions, which account for the specific elevation of predicted IBC in partners during active prosocial behavior. For naïve animals encountering an opposite-sex conspecific for the first time, the interaction is simultaneously a potential mating opportunity and a potential threat, generating heightened attentional engagement across the full range of social behaviors, spanning pro- and anti-social spectra, and producing broadly elevated IBC. After bonding, an unfamiliar individual is no longer a potential partner; instead, they become a potential threat both to the animal’s physical safety and to the integrity of the established pair bond. This shifts the salience of the interactions toward agonistic encounters, which may explain the elevated IBC specifically during antisocial behavior in stranger pairs after bonding. This framework is supported by the predictions of our Bayesian model and offers testable hypotheses for future work examining the neural substrates of these distinct IBC-generating mechanisms.

Several potential limitations merit consideration. First, partner preference scores provided a useful index of bond strength but reflect a single behavioral assay assessed at a single time point; future work with richer longitudinal behavioral measures of bond quality would strengthen inferences about IBC and its relation to bond strength. Second, our recordings were confined to the mPFC, but previous work has shown correlated c-Fos activity between prairie vole partners across the PFC, amygdala, and preoptic area and ventromedial hypothalamus^48^. As c-Fos is an immediate early gene, its correlated expression may represent real-time correlated neural activity. Extending hyperscanning approaches to these regions, and other subcortical regions necessary for pair bonding, including the nucleus accumbens and lateral septum, will be essential for a mechanistic account of IBC in social bonding^48–51^. Finally, the present study cannot fully resolve whether IBC is a cause or consequence of bond formation and bond strength. Establishing the functional significance of IBC and its relationship to the subjective experience of social attachment represents a central goal for future investigation. Prairie voles’ amenability to chemogenetic and optogenetic approaches, combined with the methodological framework established here, provides a tractable path toward addressing these questions.

Together, our findings suggest that IBC does not reflect a single social process but rather a constellation of behavioral, cognitive, and affective influences whose contributions depend on the social and relational context of the interaction. This agrees with previous hypotheses on the underpinnings of neural synchrony^51^, now explicitly evaluated longitudinally in the context of pair bonding. This also underscores the importance of studying complex social processes in preclinical models that exhibit social attachment, building substantively on and synthesizing fundamental discoveries across humans, mice, and bats.

## Acknowledgements

We thank Greyson Seale and Anna McTigue for assisting in the analyses used in Figures 1 and 3, as well as all members of the Donaldson lab for useful discussions. We would also like to thank Patrick Mahoney for his help editing the manuscript and Erin Murphy for her assistance in designing and validating the statistical models in Figures 2 and 4. This work was supported by NIH U01NS131406 (Z.R.D. and Y.K.), F31MH130143 (K.M.).

## Methods

### Resource Availability

#### Lead contacts

Further information and requests for resources and reagents should be directed to and will be fulfilled by the lead contacts, Zoe R. Donaldson and Kathleen Z. Murphy.

#### Materials availability

Behavior testing chamber designs can be accessed at https://github.com/donaldsonlab/PPT-Chamber.

#### Data and code availability

Data used in each figure panel have been deposited on GitHub: https://github.com/donaldsonlab/Interbrain_correlation and are publicly available as of the date of publication. Any additional information required to reanalyze the data reported in this paper is available from the lead contacts upon request.

### Experimental Method details

#### Animals

Prairie voles were bred in-house. Breeders were sourced from colonies at Cornell University, Emory University, and UC Davis. All lineages are descendants of wild animals captured in Illinois. Colony rooms were kept at 23-26°C on a 14:10 light:dark cycle. All procedures occurred during the light phase. Voles had ad libitum access to water and rabbit chow (5326-3 by PMI Lab Diet). Rabbit chow was supplemented with sunflower seeds, dehydrated fruit bits, and alfalfa cubes. Home cages were enriched with cotton nestlets, plastic houses, and PVC tubes. At postnatal day 21 (p21), animals were weaned and placed into standard static rodent cages (17.5 l. x 9.0 w. x 6.0 h. in.) at a density of 2-4 same-sex prairie voles. Voles underwent surgery for viral injections and implants between p60 and p100 and returned to their same-sex cohousing cages. Upon opposite-sex pairing, if there were signs of aggression, voles were placed into a clean standard static rodent cage with a perforated acrylic divider for 3 days, then moved into smaller static rodent cages (11.0 l. x 6.5 w. x 5.0 h. inches) for the remainder of the experiment. All procedures were approved by the University of Colorado Institutional Animal Care and Use Committee.

#### AAV infusion and ferrule implantation

Experimental voles underwent viral infusion and ferrule implantation surgery between p60 and p100. Anesthesia was induced using 4% isoflurane at an oxygen flow rate of 1L/min in an induction chamber. Throughout surgery, anesthesia was maintained with 1–3% isoflurane at an oxygen flow rate of 1L/min in a head-fixed stereotactic frame (David Kopf, Tujunga, CA). Body temperature was maintained at 37°C using a closed-loop heating pad with a rectal thermometer (David Kopf, Tujunga, CA). Eyes were lubricated with ophthalmic ointment (Puralube Vet Ointment). The scalp was shaved and disinfected with 70% ethanol and betadine. About 1.5 cm^2^ of skin was removed over bregma. One 0.5 mm guide hole was drilled in each parietal plate, and anchoring screws were inserted. The head was leveled in the anterior-posterior plane, and a 0.5 mm hole was drilled at + 2 mm AP and +0.4 mm ML from Bregma. A Nanoject syringe (Drummond Scientific, Broomall, PA) was used to inject 200 nL of virus at a rate of 1nL/sec at −2.7, −2.6, and −2.5 mm DV. We used AAV1-hSyn-GCaMP6f-WPRE.SV40 from Addgene (100837-AAV1) at a titer of 6.5 ×10^12^ GC/ml. A fiber-optic ferrule (0.2 mm diameter, 3.5 mm long; Neurophotometrics) was slowly lowered to a depth of 2.5 mm DV. The ferrule and screws were affixed to the skull with Loctite 454 and cured with liquid dental cement. The initial layer was covered with additional Loctite with black carbon powder (Sigma). Sustained-release meloxicam (2 mg/kg), enrofloxacin (5 mg/kg), and saline (up to 2 mL) were administered subcutaneously perioperatively to prevent pain, infection, and dehydration, respectively. Additionally, enrofloxacin (5 mg/kg) and saline (1 mL) were administered subcutaneously for the three days following surgery. Experiments started at least 21 days following surgery to allow for recovery and optimal viral expression. Ferrule placement and viral expression were confirmed posthumously.

#### Histology

Upon completion of all experimental sessions, voles were transcardially perfused with 4% paraformaldehyde in phosphate-buffered saline. The head was removed and post-fixed for 24 hours in 4% paraformaldehyde before extracting the brain. The brain was equilibrated in 30% sucrose, sectioned in 50 μm slices using a sliding freezing microtome (Leica), and mounted on slides. Slides were imaged on an Olympus IX83 slide-scanning fluorescence microscope. Viral and ferrule location were confirmed using AP histology (https://github.com/petersaj/AP_histology). Four male voles were excluded due to off-target implant placement or suboptimal expression.

#### IBC recording paradigm

Voles were briefly anesthetized with isoflurane (<30 sec) to allow experimenters to attach fiber photometry patch cables. They were placed on either side of an opaque divider in a two-chamber plexiglass arena (50.0 cm long, 20.0 cm wide, and 30.0 cm tall). The experiment started once both subjects had fully recovered and were ambulating normally. After 10 minutes of recording, the divider was removed, and the subjects were allowed to interact freely. All interactions were monitored to prevent patch cables from tangling and prevent excessive aggression. One interaction period was aborted early, due to repeated bouts of tumble fighting lasting over 5 minutes; this recording was excluded from analyses. After 20 minutes, the voles were separated, and the divider was put back in place. Recording continued for another 10 minutes. After recording, voles were briefly anesthetized to remove the patch cables and returned to their home cage. Each animal was recorded twice per recording day: once interacting with their partner and once with an unfamiliar vole. The order of recordings was counterbalanced between animals. Behavior was captured with an overhead camera. Recordings were done on two pairs simultaneously. Opaque arena walls prevented pairs recorded simultaneously from seeing each other.

#### Fiber photometry Data acquisition

Fluorescence was simultaneously acquired from voles using the Neurophotometrics (NPM) V2 system with 200 µM core optical fibers purchased from Neurophotometrics and 4-way branching cables from Doric Lenses Inc. Light was delivered at a frame rate of 60 fps, alternating between 470 nm and 415 nm. The LED power for each wavelength was set to 50 µW at the optical fiber tip. Overhead video was collected at 30 fps. Fiber photometry data and video were collected using Bonsai^52^, and each frame was marked with a computer timestamp, which was then used to align fiber photometry data to the behavior video data.

#### Partner preference test

Partner preference tests were performed as described in Scribner et al. 2020^53^. Both partner and novel animals were tethered to the end walls of three-chamber plexiglass arenas (76.0 cm long, 20.0 cm wide, and 30.0 cm tall). Tethers consisted of an eye bolt attached to a chain of fishing swivels that slid into the arena wall. Animals were briefly anesthetized with isoflurane and attached to the tether using a zip tie. Three pellets of rabbit chow and water bottles were accessible to each tethered animal. After tethering the partner and novel animals, experimental animals were placed in the center chamber of the arena. At the start of the test, the opaque dividers between the chambers were removed, allowing the subject to move freely about the arena for 3 hours. Overhead cameras (Panasonic WVCP304) were used to record 8 tests simultaneously. Two partner preference tests were run for each pair of voles within a day: one with the female as the test animal and one with the male as the test animal. The order of the tests was counterbalanced between pairs. Novel animals were sexually naïve opposite sex conspecifics that did not undergo surgical procedures.

### Quantification and statistical analysis

#### Fiber-photometry data pre-processing

Fiber photometry data were analyzed using our previously published pipeline, PhAT^54^. To preprocess the data, we corrected for photobleaching in the 470 and 415 channels by fitting and subtracting a bi-exponential function. The 415 nm channel was then used as a reference to correct for motion artifacts in the 470 nm signal using TMAC^55^. Large motion artifacts (>5x signal) were identified, and these time segments were removed before the remainder of the signal was preprocessed again. One recording was excluded due to a large unidentified artifact (∼10x signal) in the 470 nm channel, but not the isosbestic channel.

#### Session-level IBC calculations and statistical analysis

The beginning and end of the Together and Separate periods were marked using BORIS and then aligned to the fiber photometry signal in PhAT. Full session IBC was computed as the Pearson’s correlation coefficient between animals’ normalized green fluorescence signals over the time periods of interest: the Together portion or the combined Separated portions. Every IBC calculation was statistically significant (p < 1.0*e*^−5^). A Fisher’s Z-transformation was used to normalize IBC Pearson’s R coefficients for all statistical tests^56^. All statistical tests, besides the linear and Bayesian models, were computed using the SciPy Stats, version 1.13.1, in Python^57^.

#### IBC recording distance tracking

The movement of both animals in each test was scored using TopScan High-Throughput software v3 (CleverSys Inc). The distance between voles was calculated as the Euclidean distance between their centroids, as identified by TopScan. The data were smoothed using a 1-second moving average. Frames where the distance between voles exceeded the chamber length were excluded as potential tracking errors. Because tracking sometimes failed to provide meaningful inter-animal distance during contact (centroids collapsed and distance dropped to 0 cm), any periods where distances below 5 cm were increased to 5 cm to indicate contact.

#### Partner preference test analysis

The movement of all three animals in each test was scored using TopScan High-Throughput software v3.0 (CleverSys Inc). Huddling was operationalized as immobile social contact, using the parameters identified in Ahern et al. 2009^58^. The partner preference score was calculated as (partner huddle time/[partner huddle time + novel huddle time]) × 100%. Animals with partner preference scores below 50% (novel preferring animals) were considered non-bonders. Any post-bonding IBC score that included a non-bonder was excluded from statistical comparisons between group means, as they did not meet the criteria to be considered bonded partners. However, they were not excluded from measures that looked for correlations between bond strength and IBC to allow for a full examination of the individual differences between bond strength and IBC in paired animals.

#### Linear mixed model

We used a Linear Mixed-Model to interpret the contributions of distance, time in the session, relationship, and bond strength on IBC throughout a session. The recording session was included as a random intercept to account for any differences in baseline IBC due to viral expression, cable connection, LED power, etc. IBC was calculated using a 120-sec sliding window with a 30-sec step size. Within each window, IBC was quantified as the Pearson correlation coefficient between normalized fluorescence traces for each animal and then transformed using Fisher’s *z* transformation. The relationship between the animals was included as a categorical fixed effect. In the first model, the sexually naive group was split into two groups: naive 1 and naive 2 (future partners), to account for the potential of IBC between partners upon first meeting, and to predict bond strength after pairing. After finding no difference between naive groups, these groups collapsed into a single naïve group in the second model. In both models, bond strength was defined as the average partner preference score of the animals within each recording, distance was defined as the median Euclidean distance between animal centroids for each 120-sec window, and time was defined as time since the start of the together portion of the IBC recording. All continuous predictors were normalized to unit variance. Bond strength was centered on the mean score across recordings (81.02%), distance was centered at the mean distance across the entire dataset (17.51 cm), and time was centered at session onset (0 min). These choices allowed for reasonable comparisons of model terms between different predictors and intuitive interpretations of intercept values.

Likelihood ratio tests were used to evaluate interaction terms, sequentially removing nonsignificant terms while respecting model hierarchy. Only interaction terms that significantly improved model performance were retained in the final model. Higher-order terms were tested and excluded prior to evaluating lower-order terms. For both the 4-group model and the 3-group model (combined naïve groups), the 4^th^ order term (Distance × Bond Strength × Relationship × Time) did not significantly improve model performance (4-group model: χ²_4_ = 3.28, p = 0.513; 3-group model: χ²_3_ = 0.11, p = 0.991). The Distance × Bond Strength × Relationship interaction term also failed to significantly improve either model’s performance (4-group model: χ²_3_ = 1.62, p = 0.655; 3-group model: χ²_2_ = 0.83, p = 0.662); as such, these terms were excluded from the final models. All other three-way interaction terms significantly improved both models and were retained (Distance × Bond Strength × Time: 4-group model: χ²_1_ = 5.09, p = 0.024; 3-group model: χ²_1_ = 5.58, p = 0.018; Distance × Relationship x Time: 4-group model: χ²_3_ = 14.513, p = 0.002; 3-group model: χ²_2_ = 14.73, p =0.0006; Bond Strength × Relationship × Time: 4-group model: χ²_3_ = 36.07, p = 7.5e-8; 3-group model: χ²_2_ = 35.36, p = 2.1e-8).

To test whether the models performed above chance, we conducted permutation tests. Permutations were created by randomly shuffling recording level predictors (Bond strength and Relationship) and circularly permuting time and distance using a different randomly generated shift for each recording. Both models’ log-likelihood scores were significantly larger than expected when compared to our randomly generated distribution (Supp. Fig. 2B-C; p < 0.0001, 9,999 permutations).

#### Bayesian model for behavior and IBC

A Bayesian model was used to estimate the contribution of each behavior category to IBC. Using a 120-sec sliding window with a 30-sec step, we calculated IBC using Pearson’s correlation. We used a Fisher’s Z transform on the R-value to normalize the IBC values. We then calculated the proportion of time animals spent in one of 10 behavior categories in each window, providing a value for each behavior category. We accounted for the compositional nature of behavior proportion values using a centered log ratio (CLR) transform. Unidentified behaviors, which made up only 0.43% of the dataset, were included when calculating behavior category proportions but excluded from the model. To avoid perfect collinearity, nonsocial behavior was also excluded from the model, serving as the default behavior category.

We modeled Fisher *z*-transformed IBC as a function of behavioral composition, relationship, and their interaction:

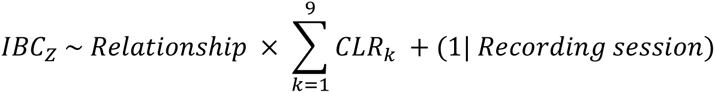

where CLR_k_ represents the standardized centered log-ratio value for each included behavioral category. A random intercept for recording session was included to account for repeated sliding windows within the same session.

The model was fit with a Gaussian likelihood and identity link, using the default weakly informative priors implemented. These priors are automatically scaled based on the data and include normal priors on regression coefficients, a normal prior on the intercept centered on the empirical mean of the outcome, and a Half-Student-t prior (ν = 4) on the residual standard deviation. Prior predictive sampling showed a reasonable range for expected IBC_Z_ values (Supp. Fig. 4A). Posterior sampling was performed using 4 Markov chains with 2,000 tuning (warm-up) iterations and 2,000 post-warmup draws per chain (8,000 total posterior samples), with a target acceptance rate of 0.95. Convergence was assessed using the rank-normalized split R^ statistic and effective sample size estimates (R^ = 1.00 all parameters). All parameters had bulk effective sample sizes exceeding 3,800, indicating adequate mixing and convergence. No divergent transitions were observed.

To evaluate whether the full model performed above chance, we conducted a permutation test using a structured null distribution. Behavioral predictors were circularly permuted within each recording to preserve temporal autocorrelation while disrupting their alignment with IBC. In addition, relationship labels were shuffled across recordings to disrupt group-level structure while maintaining within-recording dependencies.

For each permuted dataset, the full Bayesian model was re-fit using reduced sampling (500 draws per chain, 4 chains), and model performance was quantified using the summed log-likelihood and variance explained (R²) derived from posterior predictive samples. This procedure was repeated 500 times to generate null distributions for both metrics. The performance of the original model was compared to the permutation distributions, and one-tailed p-values were computed as the proportion of permuted models exceeding the observed model performance (Supp. Fig. 4C, p = 0.002).

Model effects were considered credibly different from zero when the 95% highest density interval (HDI) did not include zero. Differences between relationship groups for a given behavioral category were assessed using posterior contrasts of the corresponding interaction terms, and contrasts were considered credible when their 95% HDIs excluded zero. We used the Bambi, Arviz, and PyMC packages in Python for modeling and analyses^59–61^.

### Machine-learning-assisted Behavioral Classification

#### Pose estimation and tracking

Multi-animal DeepLabCut^37^ (DLC) was used to track the body parts of voles during two-chamber interactions. We used seven keypoints (Fig. 3B) that have been used previously to track mice in DLC^62^. All keypoints were labeled in each frame, even when the body part was occluded by another vole, themselves, or the chamber walls, to reduce null values in our final dataset. Frames were selected from multiple videos across different days, sexes, and relationships to increase generalizability. The initial frame selection was done using the built-in k-means algorithm to pick visually distinct frames. We iteratively trained DLC models using the ResNet-50 architecture on 95% of the labeled frames for 200,000 iterations, the suggested parameters for multi-animal DLC. After each iteration, poorly labeled frames were identified with an in-house Python script. Additional frames were labeled based on the areas of poorest model performance, then a new model was trained. The total process involved four iterations, and the final training set consisted of 924 frames from 32 different videos. The final model was used to label all videos. The predictions were then refined using the DLC built-in SARIMAX model with the suggested parameters to reduce jitter, estimate missing body part locations, and improve tracklets.

#### Feature Extraction

We then extracted behaviorally relevant information from the key points to use in our unsupervised classification. These features were designed to distinguish between distinct behaviors while reducing undesired clustering. Features were designed to be position-agnostic, identity-agnostic, and robust to noise. Each feature was calculated relative to the other vole, rather than dependent on global position or orientation. To remove the confound of the identity label while preserving the relationship between the features within each vole, we designated the vole more directly oriented towards the other vole as the “initiating” vole, the other was labeled the “receiving” vole (Fig. 2B). After clustering, the original identity could be reassigned to each animal to determine which animal was initiating each behavior. We avoided features that relied on flank, tail base, or spine points exclusively, due to their lower precision; instead, we opted for features based on fitted lines through these points or averages. Averages were calculated using medians instead of means to reduce the effect of outliers. Noise is also amplified when attempting to calculate time-dependent measures and increases with each derivative. Therefore, we used 10-frame windows to calculate the variance in speed and orientation as a substitute for a direct acceleration calculation. The final feature set included 4 inter-vole metrics and 5 metrics that were calculated for each vole, for a total of 14 metrics. These features were then standardized and weighted by importance before unsupervised clustering (Table 3).

#### Unsupervised clustering

The weighted features for all 1,607,271 frames were then reduced to 100,000 clusters using mini-batch k-means, with clusters ranging in size from 0 to 170 frames and a mean cluster size of 16.1 frames. The centroids of these clusters were then clustered further using Ward Agglomerative clustering^38,39^. This hierarchical approach allowed us to choose when to stop clustering certain branches of frames to get the best clustering precision for different behaviors. For instance, we could allow the algorithm to combine many nonsocial behaviors into one large cluster, while stopping the clustering process earlier for social behaviors that may require more precision, allowing us to distinguish, for instance, contact from non-contact investigation (Fig. 3I, Supp. Fig. 3A). The final set of behavioral clusters was labeled by watching clips of labeled behaviors made with an in-house python script to select clusters made up of a specific behavior. Clustering was implemented using scikit-learn and videos were made with OpenCV in python^63,64^.

