## Supplemental figures and tables for "Social attachment shapes interbrain synchrony"

### Supplementary Figures

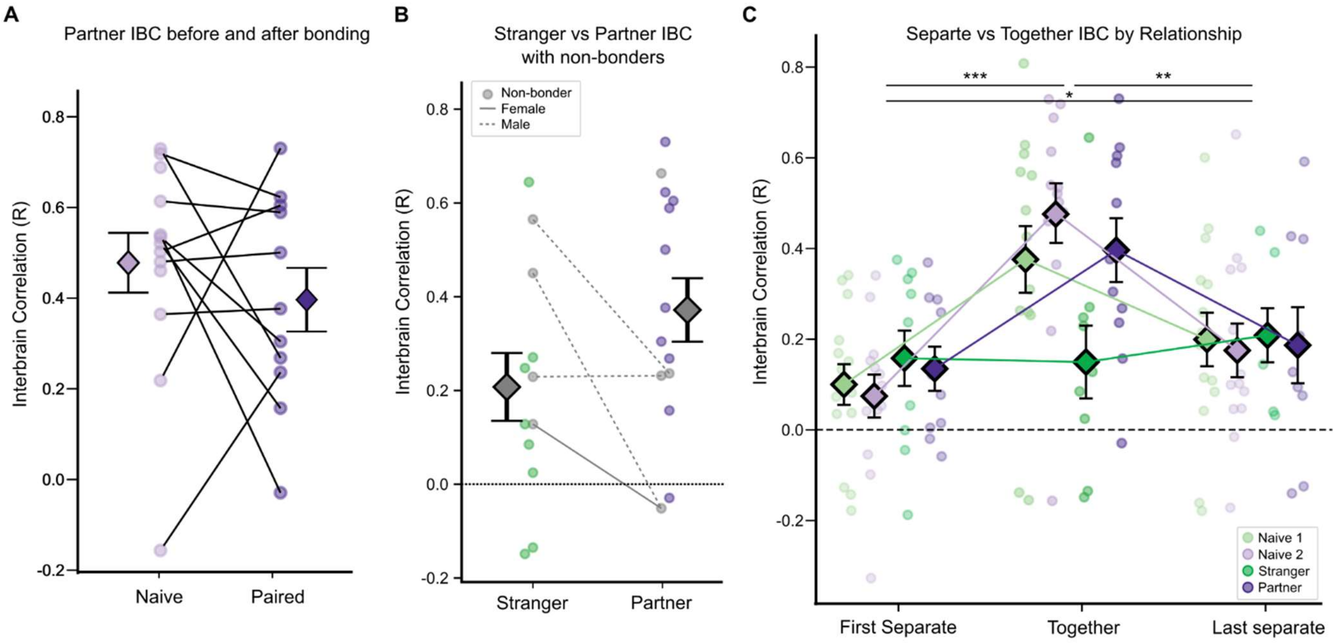


**Supplementary Figure 1. A.** IBC between partners does not increase as a function of bond formation. Lines connect scores collected from the same pair of animals, non-bonders excluded from the paired time-point (paired-samples t-test: $t_{10} = 0.657$, p =0.526) **B.** Strangers vs Partners IBC at the paired time-point with non-bonders included (Independent t-test: $t_{24} = -1.662$, p = 0.110). Lines connect Stranger and Partner sessions that had the same non-bonded animal **C.** IBC for each portion of the IBC recording by relationship, non-bonders excluded from paired groups. IBC is higher when together than either separate portion, and higher in the second separate potion than the first (Mixed-Anova: Main-effect test-portion: $F_{2,74} = 18.44$, p = 3.2e-7, Holm-corrected t-tests: $t_{40} = 5.31$, p = 1.3e-5; $t_{40} = 3.73$, p = 0.001; $t_{40} = -2.44$, p = 0.019). Test revealed no main effect of Relationship $F_{3,37} = 0.71$, p = 0.549) or Interaction effect ($F_{6,74} = 1.33$, p = 0.256). All error bars show mean and standard error. *p<0.05, **p<0.01, ***p<0.001

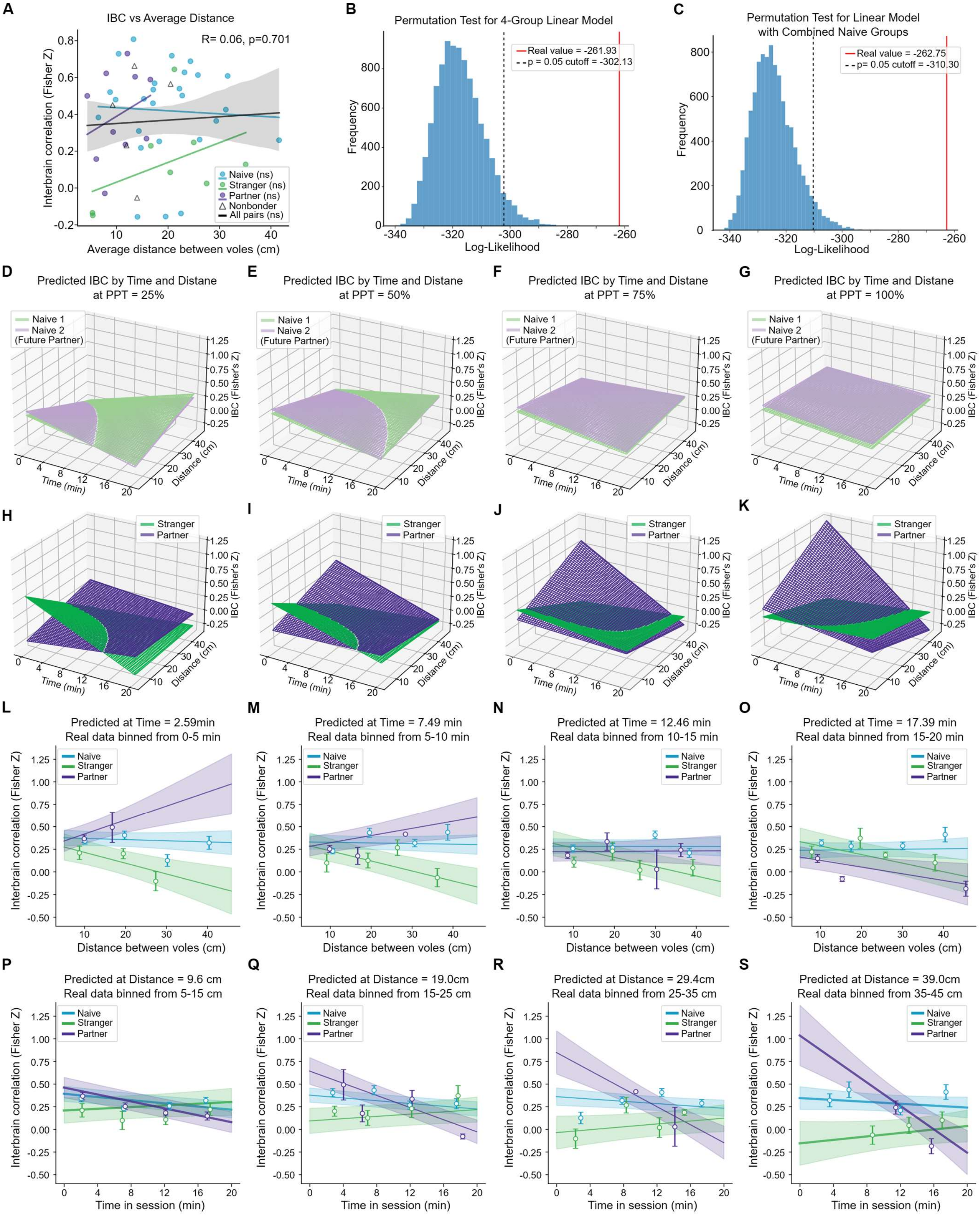

A
IBC vs Average Distance
R= 0.06, p=0.701
Interbrain correlation (Fisher Z)
Average distance between voles (cm)
Naive (ns)
Stranger (ns)
Partner (ns)
Nonbonder
All pairs (ns)
B
Permutation Test for 4-Group Linear Model
Real value = -261.93
p = 0.05 cutoff = -302.13
Frequency
Log-Likelihood
C
Permutation Test for Linear Model
with Combined Naive Groups
Real value = -262.75
p= 0.05 cutoff = -310.30
Frequency
Log-Likelihood
D
Predicted IBC by Time and Distane
at PPT = 25%
E
Predicted IBC by Time and Distane
at PPT = 50%
F
Predicted IBC by Time and Distane
at PPT = 75%
G
Predicted IBC by Time and Distane
at PPT = 100%
Naive 1
Naive 2
(Future Partner)
IBC (Fisher's Z)
Time (min)
Distance (cm)
H
I
J
K
Stranger
Partner
L
Predicted at Time = 2.59min
Real data binned from 0-5 min
M
Predicted at Time = 7.49 min
Real data binned from 5-10 min
N
Predicted at Time = 12.46 min
Real data binned from 10-15 min
O
Predicted at Time = 17.39 min
Real data binned from 15-20 min
Naive
Stranger
Partner
Interbrain correlation (Fisher Z)
Distance between voles (cm)
P
Predicted at Distance = 9.6 cm
Real data binned from 5-15 cm
Q
Predicted at Distance = 19.0cm
Real data binned from 15-25 cm
R
Predicted at Distance = 29.4cm
Real data binned from 25-35 cm
S
Predicted at Distance = 39.0cm
Real data binned from 35-45 cm
Time in session (min)

**Supplementary Figure 2. A.** Full session IBC value does not correspond to the average distance between animals during a session, either between or across groups (Pearson's correlation: all $p > 0.05$). **B.** The Linear model with all 4 relationship types preformed above chance as shown by the real Log-Likelihood value exceeding the 95% percentile of the distribution of Log-likelihoods of the models fitted on permuted data ($p<0.0001$, 9,999 permutations). **C.** The combined naïve group linear model also performed significantly above chance ($p<0.0001$, 9,999 permutations). **D-G.** Naïve groups showed very similar predicted IBC values across all distance and time values. This was true at range of bond strengths shown: (D) Stranger preference (25% PPT), (E) no preference (50% PPT), (F) partner preference (75% PPT) and (G) strong bonds (100% PPT). **H-K.** Paired groups showed a divergence in predicted IBC depending on distance and time that depends on bond strength and difference in IBC were smallest when animals showed no preference (50% PPT). **L-S.** Real binned data aligns well with linear model predictions. Only high bonding dyads (PPT > 90%) were included in real data bins (n = 30 of the 53 recordings). Each graph shows predictions at a bond strength of 97.5% PPT (mean of included recording PPT values). Real data was not adjusted to account for random intercepts included in the model. Error bars show mean and standard error. **L-O.** Show real and predicted IBC by distance for 5 minute time bins. Real data was binned again by distance for each relationship type. Lines and shaded bands show predicted IBC and 95% confidence intervals at the mean time of the real data included. **P-S.** Show real and predicted IBC by time for 10 cm distance bins. Real data was binned again by time for each relationship type. Lines and shaded bands show predicted IBC and 95% confidence intervals at the mean distance of the real data included.

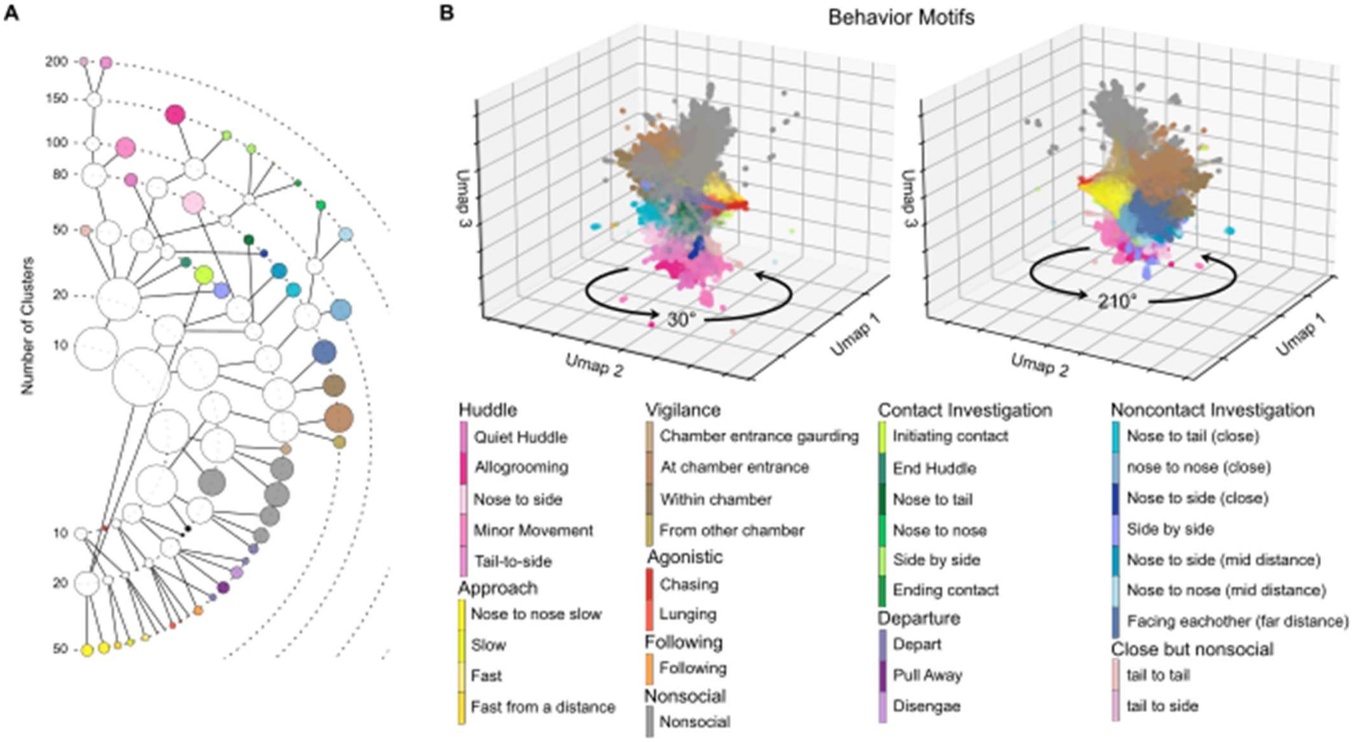


**Supplementary Figure 3. A.** Graphic showing hierarchical clustering process. The clustering process was stopped throughout at different values of clusters. Cluster that contained one behavior motif are colored accordingly. White clusters were further differentiated by identifying subclusters that contained only one behavior. **B.** Umap showing the behavior motif identified in each frame. Behavior categories correspond to expected features for those frames when compared with Figure 2F-H. Behavior motifs were sorted into 10 behavior categories.

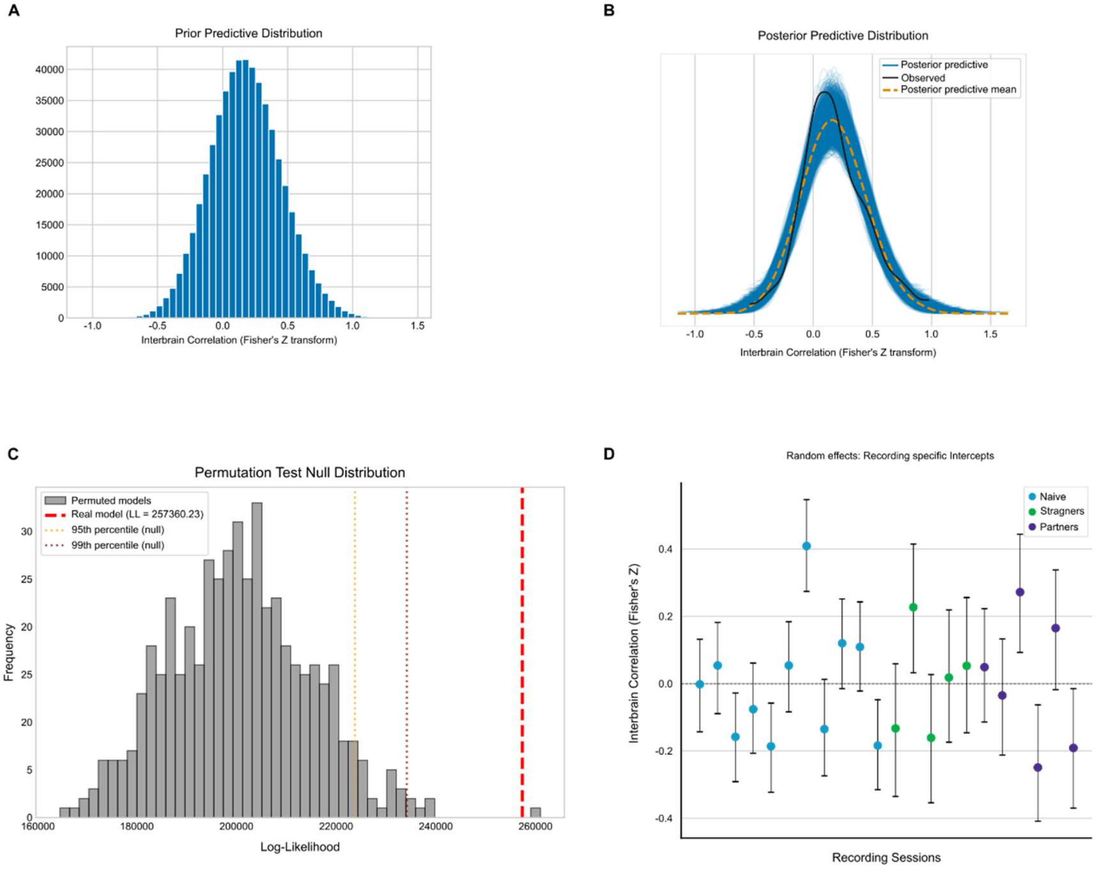


**Supplementary Figure 4. A.** Prior Predictive distribution shows default priors predict an appropriate range of IBC values. **B.** Posterior predictive distribution shows predictions are consistent with observed IBC values. **C.** The log-likelihood of our real model was significantly higher than the null distribution of log-likelihood values of models fitted on permuted data (p = 0.002, permutations = 500). **D.** Random effect intercepts for each recording colored according to the relationship between animals. Error bars represent the 95% HDI.
